# occumb: An R package for site occupancy modeling of eDNA metabarcoding data

**DOI:** 10.1101/2025.01.09.632116

**Authors:** Keiichi Fukaya, Yuta Hasebe

## Abstract

Environmental DNA (eDNA) metabarcoding has become an increasingly popular method for the rapid assessment of species distribution and diversity. However, the detection of DNA sequences of species in eDNA metabarcoding is not flawless; thus, accounting for potential false negatives (nondetection of DNA sequences of existing species) is crucial for the effective implementation of eDNA-based monitoring and research. This study introduces the occumb R package, which was developed to facilitate the easy and flexible application of community site occupancy modeling for eDNA metabarcoding that accounts for the imperfect detection of DNA sequences of species. The package provides functions for specifying models using the formula syntax, Bayesian model fitting using Markov chain Monte Carlo (MCMC), posterior inferences, and model assessments and exploring optimal study settings based on the fitted model. A case study of eDNA metabarcoding on riverine aquatic insects is illustrated using the occumb package, emphasizing the importance of ensuring biological replicates in achieving reliable species detection in eDNA metabarcoding. By making complex community site occupancy models easily accessible to users, we hope that the development of the occumb package will promote eDNA metabarcoding applications for the accurate assessment of species distribution and diversity.

## 1 Introduction

Environmental DNA (eDNA) survey is becoming more widely adopted for assessing species distribution and diversity. It can be broadly divided into the species-specific approach, which uses a primer-probe set that amplify and detect DNA sequences unique to the target species, and the metabarcoding (or amplicon sequencing) approach, which utilizes universal primer sets that amplify the DNA sequences of different species for subsequent high-throughput sequencing and taxonomic assignment (Minamoto 2022). Both approaches of eDNA surveys are generally more sensitive than conventional survey methods (Wilcox *et al*. 2016, Lugg *et al*. 2018, Bush *et al*. 2020, McColl-Gausden *et al*. 2021). However, ensuring accurate species detection remains an important issue because failure to detect DNA sequences of existing species (i.e., false negatives) may be attributed to various factors during the multistage processes of field sampling and eDNA analysis (Zhan & MacIsaac 2015, Burian *et al*. 2021). In imperfect detection, uncertainty remains when the DNA sequence of a species is undetected: the species may not be present, or it may be present but not detected. Such uncertainty can impede efficient decision-making in conservation and management.

Site occupancy models are a type of hierarchical model that deal with imperfect species detection and are widely employed in ecological studies (MacKenzie *et al*. 2017). The model includes a system submodel that describes the site occupancy of species and an observation submodel that outlines the species detection process. It allows the separate estimation of occupancy and detection probabilities, provided that repeated observations of species detection or nondetection are conducted for each site (or across an adequate number of sites). The model also includes a latent variable that indicates the existence of the species in each site, allowing for the estimation of the posterior probability that the species exists at locations where it was undetected based on Bayes’ rule.

Single-species site occupancy models have been used in species-specific eDNA applications. Given that eDNA survey follows nested sampling designs, multiscale models with a sequence of three Bernoulli (or binomial) submodels are often applied in these applications. Specifically, these models describe the existence of species at the site level, capture of the DNA sequences of species at the environmental sample level, and amplification of the DNA sequences at the polymerase chain reaction (PCR) replicate level while accounting for imperfect detection (Schmidt *et al*. 2013, Hunter *et al*. 2015, Dorazio & Erickson 2018, Stratton *et al*. 2020, McColl-Gausden *et al*. 2024). More generalized models that account for false positives in addition to false negatives have also been proposed (Guillera-Arroita *et al*. 2017, Griffin *et al*. 2020). Several R packages and software for site occupancy modeling, such as ednaoccupancy (Dorazio & Erickson 2018), msocc (Stratton *et al*. 2020), and eDNA 1.0 (Diana *et al*. 2021), have been developed for species-specific eDNA surveys.

For eDNA metabarcoding, multispecies, or community, site occupancy models have been applied. In earlier applications, metabarcoding results were aggregated to the binary form (i.e., indicating detection (1) and nondetection (0) of the DNA sequences of species from the samples) for the application of site occupancy models (Doi *et al*. 2019, Bush *et al*. 2020, McClenaghan *et al*. 2020, McColl-Gausden *et al*. 2021, Peixoto *et al*. 2023, Tetzlaff *et al*. 2024). Although such an approach can employ the traditional occupancy modeling framework with the Bernoulli-type observation submodel, it fails to explicitly describe variations in sequence read counts, which are the original data type in eDNA metabarcoding. Alternatively, Fukaya *et al*. (2022) proposed a multiscale community site occupancy model that incorporates a multinomial observation submodel for sequence reads. The proposed model facilitates a more detailed analysis of species detectability by thoroughly considering quantitative data in read counts. They also proposed a model-based approach for designing eDNA metabarcoding studies that are optimized for species detection by comparing the expected number of species detected across different study settings. These methods necessitate model fitting and prediction through a fully Bayesian approach; however, for effective application, users must prepare custom codes tailored to their problems.

Herein, we introduce the occumb R package (Fukaya 2024), which implements the community site occupancy modeling framework for eDNA metabarcoding developed by Fukaya *et al*. (2022). The package offers functionalities for performing Bayesian model fitting and posterior inferences. The package requires minimal coding work for users to build and fit their models because it allows them to specify the model form conveniently using the R formula syntax. Moreover, we present an overview of the package’s workflow and introduce a case study in which the package was used to analyze eDNA metabarcoding data of aquatic insects.

## 2 Modeling framework

### 2.1 Model formulation

This section describes the formulation of the model that the occumb package implements (Fukaya *et al*. 2022). Subsections 2.1.1 to 2.1.3 describe the most general form of the model. Subsection 2.1.4 presents a specific model example with a more restricted form.

#### 2.1.1. Observation and system models

We assume that *I* focal species detected by eDNA metabarcoding of environmental samples collected at *J* sites are modeled. At site *j* (= 1, …, *J*), *K*_*j*_ replicates of environmental samples are assumed to be collected. To ensure that the model parameters can be fully estimated, multiple replicates (i.e., *K*_*j*_ *>* 1) are required for at least a subset of the sites. We denote the sequence read counts of species for replicate *k* (= 1, …, *K*_*j*_) at site *j* obtained using high-throughput sequencing and subsequent bioinformatic processing, as ***𝒴***_*jk*_ = (*y*_1*jk*_, …, *y*_*Ijk*_).

Sequence reads are assumed to follow a multinomial distribution as shown below:

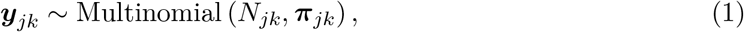

where *N*_*jk*_ = ∑_*i*_ *𝒴* _*ijk*_ is the per-sample total sequence reads, which we call sequencing depth, and ***π***_*jk*_ = (*π*_1*jk*_, …, *π*_*Ijk*_) refer to the multinomial cell probabilities expressing the relative frequency of the sequence of each species in the library of replicate *k* at site *j*. The multinomial cell probabilities are modeled as follows:

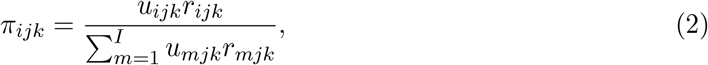

where *u*_*ijk*_ is a latent indicator variable that represents the presence (*u*_*ijk*_ = 1) or absence (*u*_*ijk*_ = 0) of the sequence of species *i* in the library of replicate *k* at site *j*, and *r*_*ijk*_ is a latent variable that is proportional to the relative frequency of the sequence of species *i*, conditional on its presence in the library of replicate *k* at site *j*. The latent indicator *u*_*ijk*_ is assumed to follow a Bernoulli distribution as shown below:

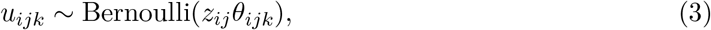

where *z*_*ij*_ is a latent indicator variable that represents the occupancy of species *i* at site *j*, and *θ*_*ijk*_ is the per replicate probability of sequence capture condition on the site occupancy of the species. The latent variable *r*_*ijk*_ is assumed to follow a gamma distribution as shown below:

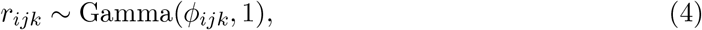

where a gamma distribution with shape *a* and rate *b* were denoted by Gamma(*a, b*). *ϕ*_*ijk*_ is a parameter that represents the relative dominance of the DNA sequence of species: species with higher *ϕ* values tend to yield more sequence reads than species with smaller *ϕ* values when their DNA sequences are present in the sequencing library as amplicons.

The latent indicator of the site occupancy of species, *z*_*ij*_, is assumed to follow a Bernoulli distribution, as shown below:

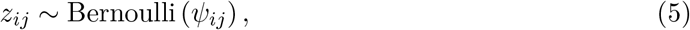

where *ψ*_*ij*_ represents the site occupancy probability of species *i* at site *j*.

Accordingly, the model contains a set of parameters, sequence relative dominance *ϕ*, probability of sequence capture *θ*, and site occupancy probability *ψ*, which relate to species detectability in different steps of eDNA metabarcoding. They can be modeled as a function of covariates by specifying a regression function similar to the generalized linear model with an appropriate link function (log for *ϕ* and logit for *θ* and *ψ*), as described in the following subsections. The occumb package allows for easy specification of covariates using the R formula syntax.

#### 2.1.2 Covariate modeling of parameters

Three types of covariates are assumed: species covariates that can take on different values for each species (e.g., traits), site covariates that can present different values for each site (e.g., habitat characteristics), and replicate covariates that can assume different values for each replicate (e.g., amount of water filtered). For each parameter, we also assume that covariates have two types of effects: effects that are species-specific and thus have index *i* and effects that are shared among all species and thus do not have index *i*. The most general form of the regression function for each parameter is expressed as follows:

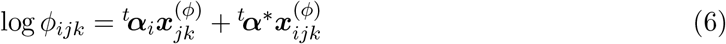

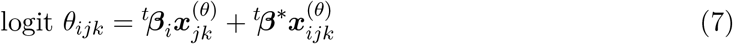

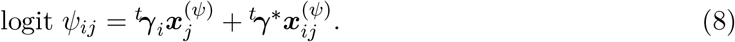

Here, 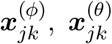 and 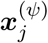 are covariate vectors for *ϕ, θ*, and *ψ*, respectively, consisting of a single one for the intercept and a set of specified covariates. ***α***_*i*_, ***β***_*i*_, and ***γ***_*i*_ are the vectors of the species-specific effects for *ϕ, θ*, and *ψ*, respectively, containing an intercept and slopes that vary between species. 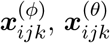 and 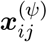 are another covariate vectors for *ϕ, θ*, and *ψ*, respectively, which are associated with the effects shared among species, ***α***^*\**^, ***β***^*\**^, and ***γ***^*\**^. The superscript *t* represents the transposition of a vector.

The covariate modeling of the parameters has some general characteristics. First, ***α***_*i*_, ***β***_*i*_, and ***γ***_*i*_ contain a species-specific intercept, whereas ***α***^*\**^, ***β***^*\**^, and ***γ***^*\**^ do not contain an intercept to ensure model parameter identifiability. Second, Equations (6)–(8) show the most general form of the model. The range of dimensions for the covariate vectors and parameters can be more limited, depending on the specified covariates; see Section 2.1.4 for specific examples of restricted forms. Third, species covariates must be associated with the shared effects because species-specific effects cannot be estimated for species covariates. Lastly, *ψ* cannot be modeled as a function of replicate covariates because it remains constant at the replicate level.

#### 2.1.3 Prior distributions

The species-specific effects are treated as random effects. A multivariate normal prior is specified for the species-specific effects as shown below:

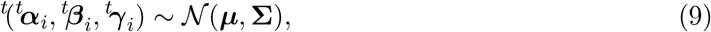

where ***µ*** and **∑** are the mean vector and covariance matrix of the species-specific effects, respectively. A vague normal prior is specified for each component of ***µ*** and each component of the shared effects ***α***^*\**^, ***β***^*\**^, and ***γ***^*\**^. A uniform prior distribution with a value greater than 0 is specified for the standard deviations of **∑**, and a uniform prior distribution that ranges from *™*1 to 1 is specified for the correlation coefficients of **∑**.

#### 2.1.4 A specific example

Fukaya *et al*. (2022) analyzed eDNA metabarcoding data of freshwater fish. They used the degree of primer-template mismatches (the total number of mismatched bases in the priming region of the forward and reverse primers) as a species covariate (*mismatch*_*i*_) and a binary indicator of the absence of aquatic and riparian vegetation at the riverbank as a site covariate (*riverbank*_*j*_). The regression model for each parameter adopted in their analysis was as follows:

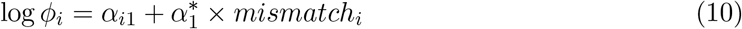

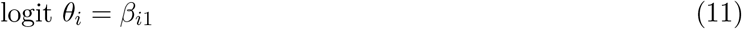

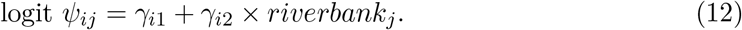

Here, *mismatch*_*i*_ is specified to have a shared effect, 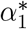 which can explain interspecific differences in the relative sequence dominance *ϕ*, and *riverbank*_*j*_ is specified to have a species-specific effect, *γ*_*i*2_, which can explain variations in site occupancy probability *ψ* for each species. Because no covariates are specified for the sequence capture probability *θ*, its value is determined only by the species-specific intercept *β*_*i*1_.

Under this specific model, the subscripts of *ϕ* and *θ* are reduced compared with Equations (6) and (7). Accordingly, in this model, Equations (3) and (4) are simplified as follows:

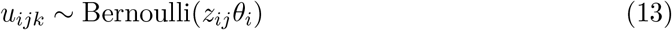

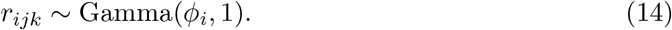

### 2.2 Model fitting and posterior inferences

To obtain posterior samples of all parameters and latent variables, the model can be fitted using a fully Bayesian approach with the Markov chain Monte Carlo (MCMC) method. The posterior samples can then be utilized to evaluate the model fit and compare the effectiveness of possible study settings regarding the assessment of species diversity.

The goodness of model fit can be evaluated using the posterior predictive check approach (Conn *et al*. 2018). The overall model fit can be inadequate or not based on the Bayesian *p*-value of an omnibus goodness-of-fit statistic, such as the Freeman-Tukey statistic.

Study settings can be compared based on the “effectiveness of species detection,” which is measured by the expected number of species detected (Fukaya *et al*. 2022). As detailed in Section 3.5, species detection can be considered effective in the assessments of local and regional species diversity levels.

## 3 Package workflow

This section briefly describes a series of data analyses supported by the package. The package’s built-in dataset on eDNA metabarcoding of freshwater fish was used to replicate the analysis performed by Fukaya *et al*. (2022). Further complementary information is available in the package vignette (https://fukayak.github.io/occumb/).

### 3.1 Building a data object

First, a data object was created using the `occumbData()` function. The `occumbData()` combines sequence read counts (`y`), species covariates (`spec_cov`), site covariates (`site_cov`), and replicate covariates `(repl_cov)` into a single object. Here, we supply the relevant elements in the built-in data `fish_raw`. Because no replicate covariate was used in the analysis, its specification is omitted, as shown below:

~~~
library(occumb)
data <- occumbData(
  y = fish_raw$y ,
  spec_cov = list(
    mismatch = fish_raw$mismatch
  ),
  site_cov = list(
  riverbank = fish_raw$riverbank
 )
)
~~~

The `y` argument takes a three-dimensional array with species, site, and replicate dimensions. In the `fish_raw` data, the dimension of `y` is 50 species, 50 sites, and three replicates. The `y` argument can accept unbalanced design data with different numbers of replicates per site, in which case zero vectors represent the data for missing replicates. Covariates are given in the named list format.

The `summary()` function can be applied to the `data` object for dataset overview.

### 3.2 Model fitting

Models are fitted using the `occumb()` function. This function generates a BUGS code inter- nally for the model specified by the R formula syntax, which is then passed to JAGS software (Plummer 2003) using the jagsUI interface (Kellner 2024) to perform a Bayesian model fitting using the MCMC method. Covariates are specified, for each parameter, through an argument identifying species-specific effects monk `(formula_phi, formula_theta, and formula_psi)` and an argument specifying effects shared across species `(formula_phi_shared, formula_theta_shared, and formula_psi_shared)`. The following code specifies the model represented by Equations (10)–(12) and stores the posterior samples in the `fit` object:

~~~
fit <- occumb(
   formula_phi_shared = ∼ mismatch ,
   formula_psi = ∼ riverbank ,
   data = data,
   parallel = TRUE
   )
~~~

The `parallel` argument sets parallel execution of the MCMC chains. The `occumb()` function also has other arguments that determine the MCMC settings (e.g., number of chains and iterations). The initial values for the MCMC chains are set randomly by default.

The `plot()` function can be applied to the `fit` object to draw a traceplot and densityplot for each parameter. A high-level summary of the MCMC results can be attained by applying `summary()` to the `fit` object.

### 3.3 Getting posterior results

Several methods can be employed to access the posterior results in the `fit` object. The `get_post_samples()` function can be used to extract the posterior samples of the parameters. For example, the following command extracts the posterior samples of psi (*ψ*) parameters from the `fit` object.

~~~
post_sample_psi <- get_post_samples(
 fit, "psi"
)
~~~

In the fitted model, *ψ* has species and site dimensions because a site covariate riverbank was specified in the `formula_psi` argument (Equation 12). Thus, `post_sample_psi` is a three-dimensional array with samples, species, and site dimensions.

The `get_post_summary()` function generates the posterior summaries of the parameters. The following command generates a posterior summary table of *ψ*, including posterior means, standard deviations, quantiles, 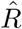 statistics, and effective sample sizes.

~~~
post_summary_psi <- get_post_summary(
  fit, "psi"
)
~~~

The prediction of *ϕ, θ*, and *ψ* values, conditional on a given set of covariate values, can be obtained using the`predict()` function. The `predict()` function has the newdata argument that accepts an optional dataset object used for prediction. If newdata is not specified, predictions are made using the covariates used for model fitting. For example, the following command generates a three-dimensional array of the posterior median and 2.5% and 97.5% quantiles as lower and upper limits, respectively, of the 95% credible interval for *ψ* of the 50 species and 50 sites, based on the values of covariates used for model fitting.

~~~
predict_psi <- predict(
  fit,
  parameter = "psi",
  type = "quantiles"
  )
~~~

For details of the parameters obtained using these functions, see the function documentation and package vignettes.

### 3.3 Assessing goodness of fit

A goodness-of-fit assessment of the model using the posterior predictive check approach can be performed by applying the `gof()` function to the `fit` object. The following command generates the value of the default statistic, the Freeman-Tukey statistic, and its Bayesian *p*-value:

~~~
gof_result <- gof(fit, cores = 4)
~~~

The cores argument specifies the number of cores used for the parallel computation of the statistic.

### 3.5 Comparing different study settings

The `eval_util_L()` and `eval_util_R()` functions estimate the expected number of species detected under specific study settings based on a model fitting result. The two functions assume species diversity assessments at different spatial scales, that is, `eval_util_L()` and `eval_util_R()` for the local and regional scales, respectively. Local assessments concentrate on the study sites used for model fitting and evaluate the average number of species anticipated to be detected at these sites based on specific combinations of the number of replicates per site and sequencing depth per replicate. For example, the following command will generate the average number of species expected to be detected at the 50 sites in the `fish` dataset when the numbe of replicates K and sequencing depth N take the values (1, 2, 3) and (1,000, 10,000, 100,000), respectively.

~~~
utilL1 <- eval_util_L(
  expand.grid(K = 1:3 ,
              N = c(1E3 , 1E4 , 1E5 )),
  fit,
  cores = 4
  )
~~~

The cores argument controls the degree of parallel computation.

If the research budget value `(budget)`, cost per sequence read for high-throughput sequencing `(lambda1)`, and cost per replicate for library preparation `(lambda2` are known, the `list_cond_L()` function can be used to obtain the set of feasible settings based on these cost and budget values. In the following commands, a list of feasible study settings (settings) is obtained using `list_cond_L()` and supplied to `eval_util_L()` to identify the optimal study setting under the specified budget and cost values:

~~~
settings <- list_cond_L(
   budget = 875000 ,
   lambda1 = 0.01 ,
   lambda2 = 5000 ,
   fit)
utilL2 <- eval_util_L(
   settings , fit,
   cores = 4
   )
~~~

The regional assessments focus on evaluating species diversity in a broader area that includes the study sites used for model fitting. The `eval_util_R()` function evaluates the total number of species expected to be detected in the region under specific combinations of the number of sites, the number of replicates per site, and sequencing depth per replicate. The `eval_util_R()` function can be used similarly to the `eval_util_L()` function. The `list_cond_R()` function is available for regional scale assessments as an analog for the `list_cond_L()` function.

## 4 A case study

Herein, we present a case study of eDNA metabarcoding of aquatic insect species in a riverine environment. Samples with different filtration water volumes were collected to estimate sequence capture probabilities (*θ*) as a function of the filtration volume. The fitted model was used to examine how increasing the filtration volume, number of biological replicates, and sequencing depth can ameliorate false negatives. The R code for the analysis is available at: https://github.com/fukayak/eDNA_aquatic_insects.

### 4.1 Materials and Methods

In July and August 2022, eDNA samples were collected from 20 sites along the Sagami River and Sakawa River in Japan. Four biological replicates with different volumes of filtered water were obtained at each site. Specifically, Sterivex HV cartridge filters (Merck, Germany) were used to filter 400, 600, 800, and 1000 ml per filter unit at each site. The MtInsects-16S universal primer set was used to amplify an mtDNA 16S rRNA region of aquatic insects (Takenaka *et al*. 2023, 2024), and the amplified DNA fragments were sequenced using an iSeq 100 sequencer (Illumina, USA). Taxonomic assignments of the representative sequences were performed up to the species level where possible using USEARCH version 11.0.667 (Edgar 2010) and BLAST version 2.13.0+ (Camacho *et al*. 2009) with the Kanagawa Prefecture DNA database for aquatic insects (https://www.pref.kanagawa.jp/docs/b4f/suigen/edna-en.html) and the NCBI GenBank database (Clark *et al*. 2016) with >99% blastn search identity. In this study, 247 taxa (hereafter referred to as ‘species’) of aquatic insects were detected, and 169 species that had occurred in four or more samples were modeled in the subsequent analysis. Three samples were omitted from which no sequence reads were obtained due to filter clogging and errors in DNA extraction. The mean and standard deviation of the sequencing depths of the remaining 77 samples were 40,416.6 and 27,795.2, respectively. More details of data acquisition are provided in Supporting Information.

The sequence read dataset was analyzed using occumb version 1.1.0. In the analysis, the order `(order)` and number of base-pair mismatch in the primer attachment site (`mismatch)` were incorporated as species covariates and filtration volume `(vol)` as a replicate covariate. The order covariate was categorical, which include six orders containing a number of aquatic insect species (Ephemeroptera, Plecoptera, Trichoptera, Odonata, Diptera, and Coleoptera), plus one “others” category of three orders (Hemiptera, Megaloptera, and Lepidoptera) for which the species detected was much fewer than other groups. The volume covariate was normalized to have a mean of zero and a variance of one. The model was fitted using the `occumb()` function, with the sequence relative dominance (*ϕ*) as a function of order and the degree of the primer-template mismatch `(formula_phi_shared = ∼ order + mismatch)` and sequence capture probability (*θ*) as a function of the order and filtration volume `(formula_theta_shared = ∼ order + vol)` , respectively. The effect of the filtered volume on *θ* was assumed to be common among species. To configure the MCMC computation, the arguments `n.iter = 250000, n.burnin = 50000,` and `n.thin = 200` were also indicated. MCMC convergence was confirmed by 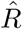 statistics for all parameters of interest being <1.1. The posterior predictive check of the fitted model revealed no significant issue of model fit (Bayesian *p*-value of the Freeman-Tukey statistic: 0.11).

The `eval_util_L()` function was used to compare the effectiveness of species detection (i.e., the average number of species expected to be detected per site) across different study settings. Two profiles of the effectiveness of species detection were obtained: the ‘unconditional’ profiles for which we set some arbitrary combinations of the number of biological replicates and sequencing depth per replicate and ‘conditional’ profiles for which we obtained a list of feasible study settings under a specific set of budget and cost values using the `list_cond_L()` function. In the latter, an estimate of the budget `(budget = 540000 JPY)` and costs `(lambda1 = 0.025` JPY and `lambda2 = 5750 JPY)` was used for the present study. The feasible combinations of the number of replicates per site and the sequencing depth per replicate were one replicate, with 850k reads, two replicates with 310k reads, three replicates with 130k reads, and four replicates with 40k reads.

### 4.2 Results

The estimated values for site occupancy probability (*ψ*), sequence capture probability (*θ*), and sequence relative dominance (*ϕ*) varied considerably between species (Fig. 1). These parameters were correlated positively (Table 1), indicating that the species with lower *ψ* values (i.e., those that would occur in a few sites) tended to be less detectable when present on a site. The estimates of the covariate effect indicated that the average of *θ* and *ϕ* varied depending on the order (Table 2). The effect of the order on *θ* was estimated to be negative, and its 95% credible interval did not overlap with zero for Coleoptera and others, indicating that the average *θ* for Coleoptera and others was lower than that for Ephemeroptera as the reference category. The effect of the order on *ϕ* was estimated to be negative, and its 95% credible interval did not overlap with zero at all levels of the factor, indicating that the average *ϕ* for Ephemeroptera was higher than for other orders. The filtered volume positively affected *θ* (Table 2), indicating that filtering more water allows for more reliable capturing of DNA sequences of species. The degree of primer-template mismatch negatively affected *ϕ* (Table 2), indicating that the species with more base mismatches at the primer binding site tend to have fewer sequence reads.

**Table 1.**
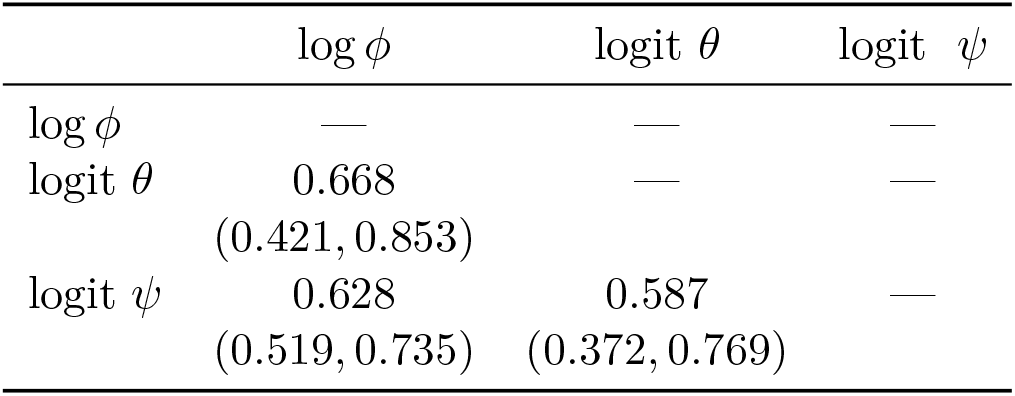
Posterior summary of the correlation coefficients between the three parameters governing the detectability of each species. The posterior median and 95% credible intervals (in brackets) are provided. The correlation coefficients were estimated as derived parameters on the link scale of the parameters, with fixed amount of the filtered volume (vol).

**Table 2.** Estimated coefficients for aquatic insects in Sagami River and Sakawa River. The coefficients for which the 95% credible interval (CI) does not overlap with 0 are highlighted in bold.

| Coefficient name given in the package | Parameter | Effect | Posterior median | Lower limit of 95% CI | Upper limit of 95% CI |
| --- | --- | --- | --- | --- | --- |
| beta_shared[1] | $\theta$ | Order (Plecoptera) | -0.251 | -0.762 | 0.412 |
| beta_shared[2] | $\theta$ | Order (Trichoptera) | 0.060 | -0.345 | 0.499 |
| beta_shared[3] | $\theta$ | Order (Odonata) | -0.520 | -1.137 | 0.250 |
| beta_shared[4] | $\theta$ | Order (Diptera) | 0.028 | -0.497 | 0.657 |
| beta_shared[5] | $\theta$ | Order (Coleoptera) | **-0.903** | **-1.446** | **-0.329** |
| beta_shared[6] | $\theta$ | Order (others) | **-0.767** | **-1.402** | **-0.009** |
| beta_shared[7] | $\theta$ | Filtered volume | **0.383** | **0.286** | **0.485** |
| alpha_shared[1] | $\phi$ | Order (Plecoptera) | **-0.707** | **-1.103** | **-0.329** |
| alpha_shared[2] | $\phi$ | Order (Trichoptera) | **-0.335** | **-0.623** | **-0.068** |
| alpha_shared[3] | $\phi$ | Order (Odonata) | **-0.659** | **-1.066** | **-0.260** |
| alpha_shared[4] | $\phi$ | Order (Diptera) | **-0.649** | **-0.976** | **-0.316** |
| alpha_shared[5] | $\phi$ | Order (Coleoptera) | **-0.549** | **-0.963** | **-0.156** |
| alpha_shared[6] | $\phi$ | Order (others) | **-0.652** | **-1.127** | **-0.189** |
| alpha_shared[7] | $\phi$ | Primer-template mismatch | **-0.161** | **-0.302** | **-0.007** |

**Fig. 1.**
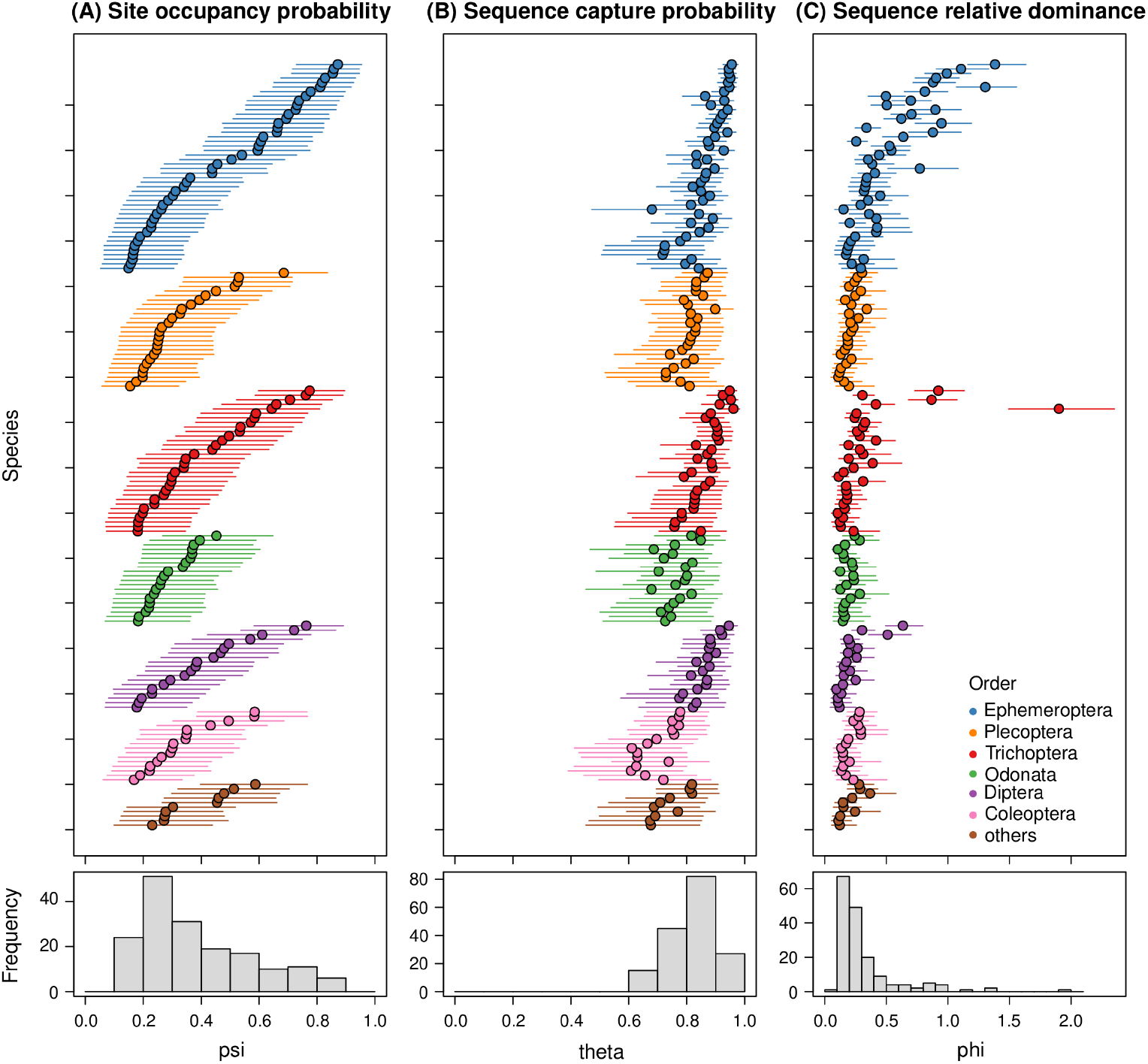
Estimates of (A) site occupancy probabilities (*ψ*), (B) sequence capture probabilities (*θ*), and (C) sequence relative dominance (*ϕ*) for aquatic insects in Sagami River and Sakawa River. In the upper panels, the posterior median and 95% credible interval for each species are represented by circles and lines, respectively. The values for *θ* are provided for a filtration volume of 1000 ml. Species are arranged according to the site occupancy probability for each order. Lower panels illustrate the histograms of the posterior medians.

The ‘unconditional’ profiles of the expected number of species detected per site indicated that the increasing filtration volume, number of biological replicates, and sequencing depth increase the expected number of species to be detected (Fig. 2). Among these three factors, the number of biological replicates exerted a particularly large effect: securing a small number of replicates was predicted to largely increase the number of species detected compared with a single replicate (Fig. 2). When comparing cases where the filtration volume and sequencing depth ‘per site’ are the same, the expected number of species detected is higher when multiple replicates are set. For example, the expected number of species detected is predicted to be 14% higher when obtaining 40k reads from each of the two replicates of 400 ml filtered (52.6 species, indicated by the filled arrow in Fig. 2) than when obtaining 80k reads from one replicate of 800 ml filtered (46.4 species, indicated by the open arrow in Fig. 2).

**Fig. 2.**
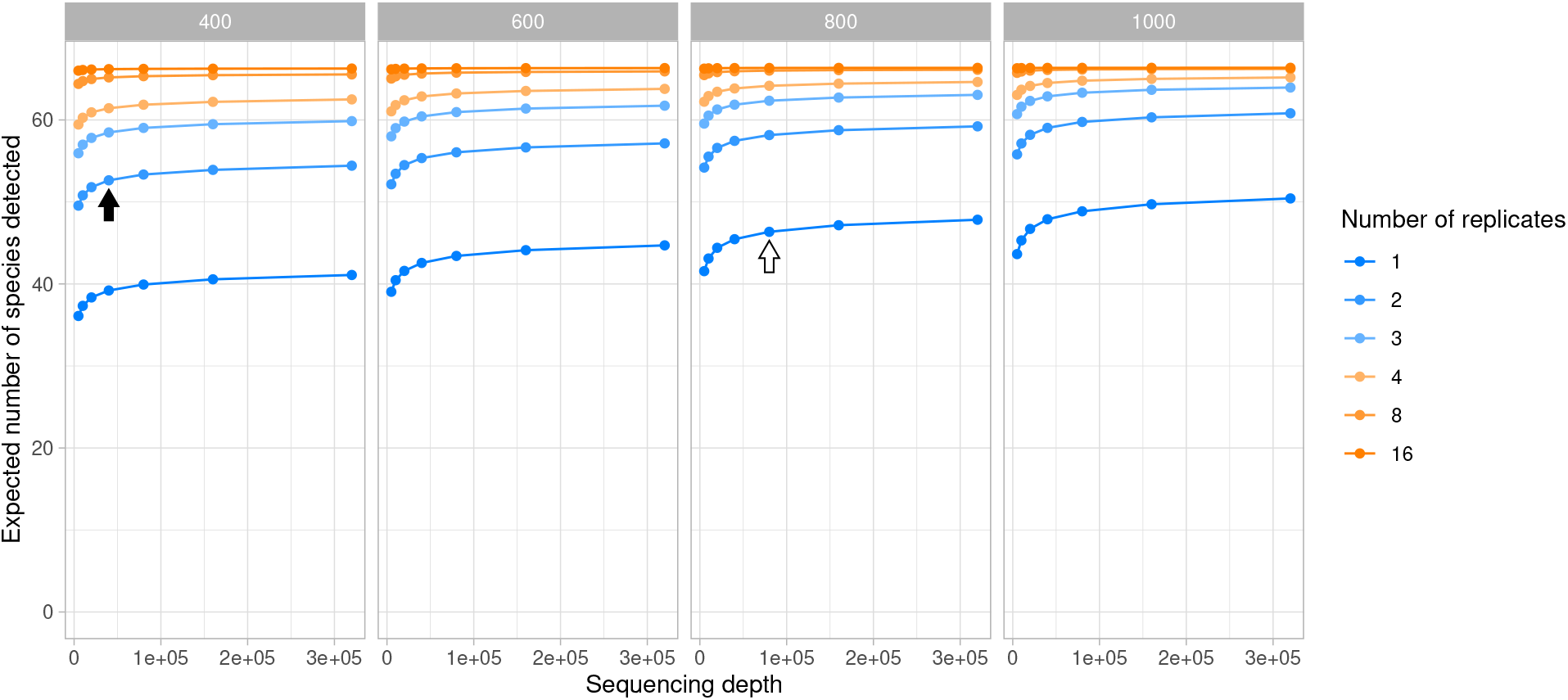
Unconditional profiles of the effectiveness of species detection of aquatic insects in Sagami River and Sakawa River. The average number of species anticipated to be detected in each site was evaluated at different levels of the filtration volume per replicate (panels), number of biological replicates per site (line colors), and sequencing depth per replicate. The title of the panels indicates the filtration volume in ml. The two conditions referred to and compared in the main text are indicated by filled (400 ml filtered, two replicates, and 40k reads) and open (800 ml filtered, one replicate, and 80k reads) arrows.

When comparing the effectiveness of species detection among the feasible study settings based on the given budget and cost values (i.e., ‘conditional’ profiles), an increase in the number of biological replicates led to an increase in the expected number of species detected (Fig. 3). The expected number of species detected is also dependent on the filtration amount. As the number of replicates increased, the expected number of species detected became less affected by changes in the filtration volume (Fig. 3), indicating that the decrease in the per-sample probability of sequence capture due to reduced filtration volume can be effectively compensated by ensuring biological replicates.

**Fig. 3.**
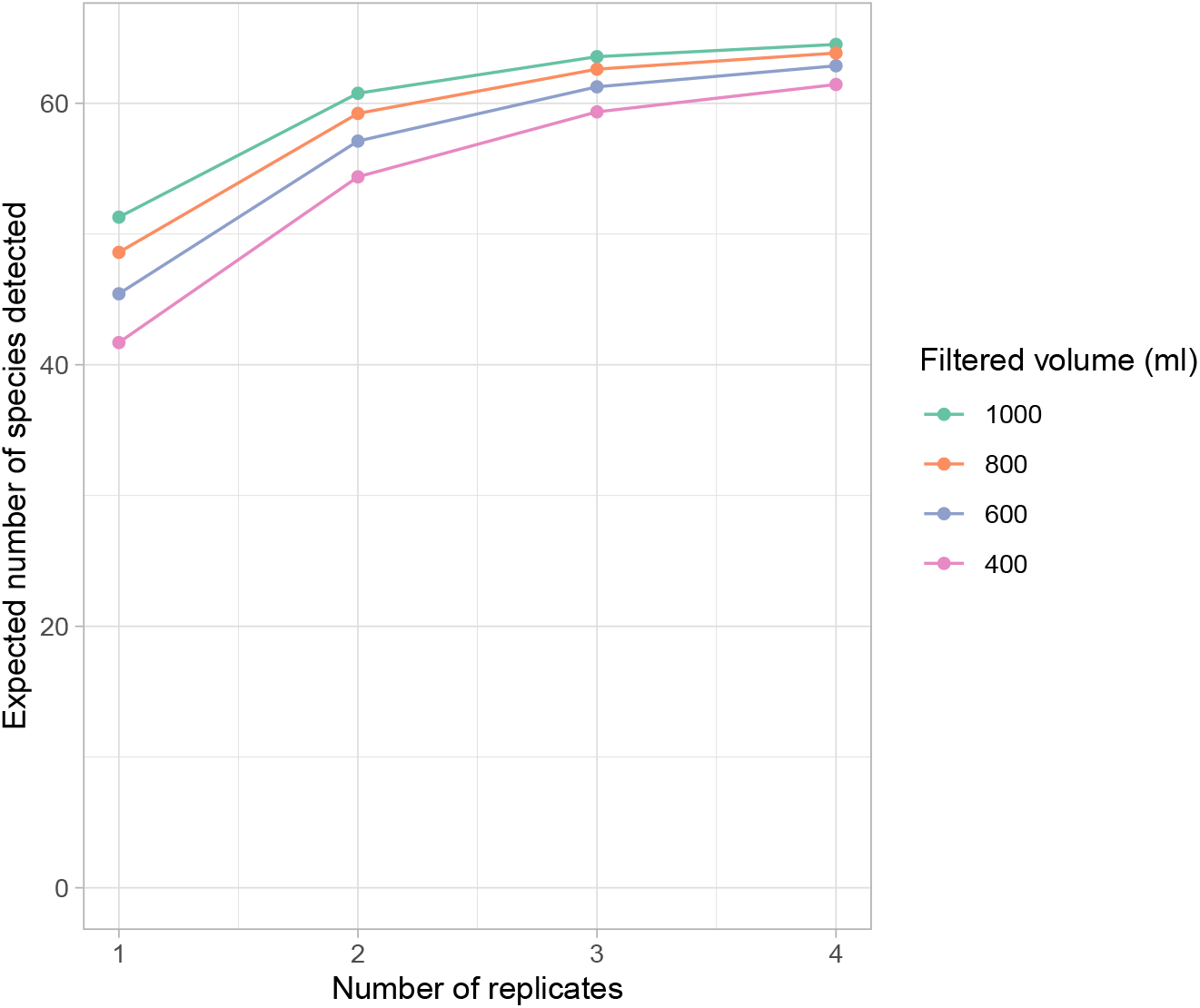
Conditional profiles of the effectiveness of species detection of aquatic insects in Sagami River and Sakawa River. The average number of species anticipated to be detected in each site was evaluated based on feasible study settings under the indicated budget and cost values.

## 5 Discussion

The occumb R package introduced herein allows users to conveniently apply the community site occupancy modeling framework for eDNA metabarcoding (Fukaya *et al*. 2022). In occumb, a variant of the community site occupancy model with a multinomial observation submodel (Fukaya *et al*. 2022) is implemented, rather than the one with a Bernoulli (or binomial) observation submodel that has been applied conventionally in eDNA metabarcoding studies (Doi *et al*. 2019, Bush *et al*. 2020, McClenaghan *et al*. 2020, McColl-Gausden *et al*. 2021, Peixoto *et al*. 2023, Tetzlaff *et al*. 2024). This feature allows the occumb package to evaluate the sources of variations in read counts and perform more detailed assessments of species detectability and study design. The functions in the package have been thoroughly unit tested and are well documented. The vignettes (https://fukayak.github.io/occumb/) of the package will complement this study by providing practical guides and references of the functions. We hope this package will facilitate eDNA metabarcoding applications for the reliable assessment of species distribution and diversity while accounting for the imperfect detection of the DNA sequences of species. The package may be applied to microbial community research and diet analysis using amplicon sequencing if the hierarchical sampling design assumed in the model is adopted.

A major challenge with the model implemented in occumb is that the MCMC algorithm may not converge quickly because of model complexity. In some applications, users may need to execute a long Markov chain until sufficient convergence is obtained. The convergence issue may be improved by data filtering, for example, by removing sequence reads with low counts from each sample or removing species that occur at extremely low frequencies, as was done in the present study of aquatic insects. Such procedures could also improve inferences by removing false-positive reads caused by PCR errors, for example, and cause biases in parameter estimation (Fukaya *et al*. 2022). The 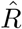 value of the latent variables *z* and *π* may not be sufficiently close to unity when its posterior probability is concentrated on zero and the chains rarely transition to nonzero values. When the 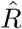 statistics of the parameters in the model proposes a convergence, except for *z* and *π*, visually checking the traceplot of *z* and *π* is recommended to ensure the absence of convergence issues. The package currently relies on JAGS (Plummer 2003) to run MCMC. The addition of an option to work with other MCMC engines that implement faster algorithms will be essential in future package development.

Applying the occumb package to the sequence read dataset of aquatic insects enables the characterization of the variations in species detectability and identify effective sampling strategies in eDNA metabarcoding. The positive correlations between the parameters governing species detectability (Table 1) implies that, in general, the ‘dominant’ species that widely exist can be detected reliably, whereas the ‘nondominant’ species that only exist in limited locations can have a higher risk of false negatives. The negative effect of `mismatch` on the relative sequence dominance *ϕ* (Table 2) indicates that a relatively smaller read count will be assigned to species with a greater degree of primer-template mismatches; thus, such species will have a higher risk of false negatives. The results confirm that the reduced PCR amplification efficiency caused by primer-template mismatches can decrease sequence reads and, therefore, species detectability in eDNA metabarcoding (Piñol *et al*. 2015, Shelton *et al*. 2016, Liu *et al*. 2023). The estimated effects of order on the sequence capture probability *θ* and the relative sequence dominance *ϕ* (Table 2) indicate that species detectability generally differs between orders. Considering that the effect of `mismatch` has been accounted for, the estimated effects of `order` might reflect, at least in part, the differences in the abundance or biomass between taxa. The positive effect of vol on the sequence capture probability *θ* (Table 2) indicates that an increase in the filtration volume can decrease false-negative results. However, in assessing the effectiveness of species detection, the importance of ensuring biological replicates was emphasized over increasing filtration volume (Figs. 2 and 3). The ‘conditional’ profiles indicated that increasing the number of replicates can mitigate the increased risk of false negatives caused by decreased filtration volume (Fig. 3). Considering that an increase in the filtration volume can diminish survey efficiency by increasing the likelihood of filter clogging (Takasaki *et al*. 2021, Peixoto *et al*. 2023), our findings indicate that ensuring multiple replicates with moderate filtration volumes per filter can be an efficient sampling strategy that balances rapid filtration processes with effective species detection.

## Acknowledgments

We thank Taku Kadoya for inspiring the authors to develop the package. We thank Koji Makiyama, Shinya Uryu, and Kentaro Matsuura for contributing to the development of package testing. We are also grateful to Mito Ikemoto and Kentaro Uehira for their valuable feedback through a package trial and a documentation review. Mika Takahashi contributed to the recent development of the package. Funding was provided by Japan Society for the Promotion of Science (KAKENHI, 20K06102 and 23H02240) and Japan Science and Technology Agency (COI-NEXT, JPMJPF2206) to KF.

## References

Burian, A., Mauvisseau, Q., Bulling, M., Domisch, S., Qian, S. & Sweet, M. (2021) Improving the reliability of eDNA data interpretation. Molecular Ecology Resources, 21, 1422–1433.

Bush, A., Monk, W.A., Compson, Z.G., Peters, D.L., Porter, T.M., Shokralla, S., Wright, M.T., Hajibabaei, M. & Baird, D.J. (2020) DNA metabarcoding reveals metacommunity dynamics in a threatened boreal wetland wilderness. Proceedings of the National Academy of Sciences, 117, 8539–8545.

Camacho, C., Coulouris, G., Avagyan, V., Ma, N., Papadopoulos, J., Bealer, K. & Madden, T.L. (2009) Blast+: architecture and applications. BMC Bioinformatics, 10, 421.

Clark, K., Karsch-Mizrachi, I., Lipman, D.J., Ostell, J. & Sayers, E.W. (2016) GenBank. Nucleic Acids Research, 44, D67–D72.

Conn, P.B., Johnson, D.S., Williams, P.J., Melin, S.R. & Hooten, M.B. (2018) A guide to Bayesian model checking for ecologists. Ecological Monographs, 88, 526–542.

Diana, A., Matechou, E., Griffin, J.E., Buxton, A.S. & Griffiths, R.A. (2021) An RShiny app for modelling environmental DNA data: accounting for false positive and false negative observation error. Ecography, 44, 1838–1844.

Doi, H., Fukaya, K., Oka, S.i., Sato, K., Kondoh, M. & Miya, M. (2019) Evaluation of detection probabilities at the water-filtering and initial PCR steps in environmental DNA metabarcoding using a multispecies site occupancy model. Scientific Reports, 9, 3581.

Dorazio, R.M. & Erickson, R.A. (2018) ednaoccupancy: An R package for multiscale occupancy modelling of environmental DNA data. Molecular Ecology Resources, 18, 368–380.

Edgar, R.C. (2010) Search and clustering orders of magnitude faster than BLAST. Bioinformatics, 26, 2460–2461.

Fukaya, K. (2024) occumb: Site Occupancy Modeling for Environmental DNA Metabarcoding. R package version 1.1.0.

Fukaya, K., Kondo, N.I., Matsuzaki, S.i.S. & Kadoya, T. (2022) Multispecies site occupancy modelling and study design for spatially replicated environmental DNA metabarcoding. Methods in Ecology and Evolution, 13, 183–193.

Griffin, J.E., Matechou, E., Buxton, A.S., Bormpoudakis, D. & Griffiths, R.A. (2020) Modelling environmental DNA data; Bayesian variable selection accounting for false positive and false negative errors. Journal of the Royal Statistical Society Series C: Applied Statistics, 69, 377–392.

Guillera-Arroita, G., Lahoz-Monfort, J.J., van Rooyen, A.R., Weeks, A.R. & Tingley, R. (2017) Dealing with false-positive and false-negative errors about species occurrence at multiple levels. Methods in Ecology and Evolution, 8, 1081–1091.

Hunter, M.E., Oyler-McCance, S.J., Dorazio, R.M., Fike, J.A., Smith, B.J., Hunter, C.T., Reed, R.N. & Hart, K.M. (2015) Environmental DNA (eDNA) sampling improves occurrence and detection estimates of invasive Burmese pythons. PLoS ONE, 10, e0121655.

Kellner, K. (2024) jagsUI: A Wrapper Around ‘rjags’ to Streamline ‘JAGS’ Analyses. R package version 1.6.2.

Liu, M., Burridge, C.P., Clarke, L.J., Baker, S.C. & Jordan, G.J. (2023) Does phylogeny explain bias in quantitative DNA metabarcoding? Metabarcoding and Metagenomics, 7, e101266.

Lugg, W.H., Griffiths, J., van Rooyen, A.R., Weeks, A.R. & Tingley, R. (2018) Optimal survey designs for environmental DNA sampling. Methods in Ecology and Evolution, 9, 1049–1059.

MacKenzie, D.I., Nichols, J.D., Royle, J.A., Pollock, K.H., Bailey, L. & Hines, J.E. (2017) Occupancy Estimation and Modeling: Inferring Patterns and Dynamics of Species Occurrence. Elsevier.

McClenaghan, B., Compson, Z.G. & Hajibabaei, M. (2020) Validating metabarcoding-based biodiversity assessments with multi-species occupancy models: a case study using coastal marine eDNA. PLoS ONE, 15, e0224119.

McColl-Gausden, E.F., Griffiths, J., Weeks, A.R. & Tingley, R. (2024) Using eDNA sampling to identify correlates of species occupancy across broad spatial scales. Diversity and Distributions, 30, e13926.

McColl-Gausden, E.F., Weeks, A.R., Coleman, R.A., Robinson, K.L., Song, S., Raadik, T.A. & Tingley, R. (2021) Multispecies models reveal that eDNA metabarcoding is more sensitive than backpack electrofishing for conducting fish surveys in freshwater streams. Molecular Ecology, 30, 3111–3126.

Minamoto, T. (2022) Environmental DNA analysis for macro-organisms: species distribution and more. DNA Research, 29, 1–9.

Peixoto, S., Mota-Ferreira, M., Chaves, C., Velo-Antón, G., Beja, P. & Egeter, B. (2023) Multi-species occupancy modeling reveals methodological and environmental effects on eDNA detection of amphibians in temporary ponds. Environmental DNA, 5, 796–811.

Piñol, J., Mir, G., Gomez-Polo, P. & Agustí, N. (2015) Universal and blocking primer mismatches limit the use of high-throughput DNA sequencing for the quantitative metabarcoding of arthropods. Molecular Ecology Resources, 15, 819–830.

Plummer, M. (2003) JAGS: a program for analysis of Bayesian graphical models using Gibbs sampling. Proceedings of the 3rd International Workshop on Distributed Statistical Computing, volume 124, pp. 1–10. Technische Universit at Wien, Austria.

Schmidt, B.R., Kéry, M., Ursenbacher, S., Hyman, O.J. & Collins, J.P. (2013) Site occupancy models in the analysis of environmental DNA presence/absence surveys: a case study of an emerging amphibian pathogen. Methods in Ecology and Evolution, 4, 646–653.

Shelton, A.O., O’Donnell, J.L., Samhouri, J.F., Lowell, N., Williams, G.D. & Kelly, R.P. (2016) A framework for inferring biological communities from environmental DNA. Ecological Applications, 26, 1645–1659.

Stratton, C., Sepulveda, A.J. & Hoegh, A. (2020) msocc: Fit and analyse computationally efficient multi-scale occupancy models in R. Methods in Ecology and Evolution, 11, 1113–1120.

Takasaki, K., Aihara, H., Imanaka, T., Matsudaira, T., Tsukahara, K., Usui, A., Osaki, S. & Doi, H. (2021) Water pre-filtration methods to improve environmental DNA detection by real-time PCR and metabarcoding. PLoS ONE, 16, e0250162.

Takenaka, M., Hasebe, Y., Yano, K., Okamoto, S., Tojo, K., Seki, M., Sekiguchi, S., Jitsumasa, T., Morohashi, N., Handa, Y. & Sakaba, T. (2024) Environmental DNA metabarcoding on aquatic insects: comparing the primer sets of MtInsects-16S based on the mtDNA 16S and general marker based on the mtDNA COI region. Environmental DNA, 6, e588.

Takenaka, M., Yano, K., Suzuki, T. & Tojo, K. (2023) Development of novel PCR primer sets for DNA barcoding of aquatic insects, and the discovery of some cryptic species. Limnology, 24, 121–136.

Tetzlaff, S.J., Katz, A.D., Johnson, M.D. & Sperry, J.H. (2024) Community ecology in a bottle: Leveraging eDNA metabarcoding data to predict occupancy of co-occurring species. Environmental DNA, 6, e579.

Wilcox, T.M., McKelvey, K.S., Young, M.K., Sepulveda, A.J., Shepard, B.B., Jane, S.F., Whiteley, A.R., Lowe, W.H. & Schwartz, M.K. (2016) Understanding environmental DNA detection probabilities: A case study using a stream-dwelling char Salvelinus fontinalis. Biological Conservation, 194, 209–216.

Zhan, A. & MacIsaac, H.J. (2015) Rare biosphere exploration using high-throughput sequencing: research progress and perspectives. Conservation Genetics, 16, 513–522.

