## Supporting Information for "occumb: An R package for site occupancy modeling of eDNA metabarcoding data"

### Appendix S1. eDNA metabarcoding of aquatic insect communities

#### Sampling sites and period

eDNA samples were collected from 20 sites in the Sagami (12 sites) and Sakawa (8 sites) river systems in Kanagawa Prefecture, Japan, for 6 days between July 25, 2022, and August 4, 2022. The sampling sites were selected to cover the upper and lower reaches of the rivers.

#### Water sampling and filtration

Four biological replicates were obtained at each site with varying amounts of filtered water. Water was collected from the surface of the river using new polypropylene bottles and immediately used a tubing pump (AS ONE, Japan) and Sterivex HV cartridge filters (Merck, Germany) to filter 400, 600, 800, and 1000 ml per filter unit at each site. Disposable instruments were used for parts in direct contact with the water sample during filtration and were replaced at each site to prevent contamination. On a site, the filter clogged when 600 ml of water was filtered, and we decided not to obtain replicates of 800 and 1000 ml. After filtration, the cartridges were filled with 2 ml of RNAlater Stabilization Solution (Thermo Fisher Scientific, USA), transported to the laboratory in a cooler, and stored at −20 °C until DNA extraction. On the last survey day, 1000 ml of ultrapure water was filtered using a Sterivex HV filter as a negative control for field sampling.

#### DNA extraction

DNA extraction was performed following the method of Wong *et al.* (2020). RNAlater was removed from the cartridge by injecting air from the inlet with a 50-ml syringe (Terumo, Japan). Then, a lysis buffer mix (PBS 990 µl, Buffer AL 910 µl, Proteinase K 100 µl; 2 ml of total volume) was introduced to the cartridge through the inlet and incubated at 56 °C for 30 min after sealing the both ends of the cartridge. The lysis buffer mix was collected from the cartridge by aspirating from the inlet using a 50-ml syringe and added 1 ml of molecular-grade ethanol (99.5%, Fujifilm Wako Pure Chemical Corporation, Japan) to the lysis buffer mix. Then, the mixture was purified using a DNeasy Blood and Tissue Kit (Qiagen, Germany). Adsorbed DNA was eluted with 150 µl of AE buffer to obtain a template for polymerase chain reaction (PCR). As a negative control for DNA extraction, a blank Sterivex HV filter filled with the lysis buffer mix was also prepared.

#### Library preparation

The resulting DNA extracts were subjected to library preparation for high-throughput sequencing using MtInsects-16S primers targeting the 16S rRNA region of mitochondrial DNA (Takenaka *et al.* 2023, 2024). Library preparation was performed using the double PCR approach to reduce bias in PCR amplification caused by indexed primers (O'Donnell *et al.* 2016).

The first PCR was performed in four replicates for each sample to minimize PCR dropout. The total volume of the first PCR reaction was 15 µl, containing 0.45 µl of 10 µM primer mix, 2 µl of DNA template, and 7.5 µl of KOD One PCR Master Mix (TOYOBO, Japan). The PCR was run at 26 cycles of 98 °C for 10 s, 55 °C for 5 s, and 68 °C for 5 s in 8-row tubes with individual lids to prevent contamination between samples. Four replicates of PCR products were pooled and used for the second PCR without purification. As a negative control for the PCR process, a blank PCR reaction was also prepared, and the first PCR was performed in the same manner.

The second PCR was performed in a final volume of 100 µl containing 3 µl of 10 µM primer mix, 4 µl of the unpurified first PCR product, and 50 µl of KOD One PCR Master Mix. The cycling condition of the second PCR was the same as for the first PCR; however, the cycle count was set to 10. Moreover, 40 µl of the second PCR product was subjected to the 0.6–0.8× right-side size selection workflow using SPRIselect (Beckman Coulter, USA) for purification and removal of nontargeted amplifications. The purified samples were diluted to 100 pg/µl each, after measuring the DNA concentration of the target region on a 4150 TapeStation System (Agilent, USA), and used for the third PCR. For negative controls, dilutions were made at the average dilution rate of the samples.

The third PCR was performed in a final volume of 30 µl containing 3.6 µl of 2.5 µM primer mix, 2 µl of the purified and diluted second PCR product, and 15 µl of KOD One PCR Master Mix. The PCR was run at 10 cycles of 98 °C for 10 s, 60 °C for 5 s, and 68 °C for 5 s. After equal amounts of each third PCR product were pooled, 40 µl of the mixed products were subjected to the 0.6–0.8× right-side size selection workflow using SPRIselect and stored below −20 °C until further processing.

**High-throughput sequencing**

The library was sequenced in iSeq 100 (Illumina, USA) using the iSeq 100 i1 Reagent v2 (300-cycle) with 20% PhiX (Illumina). All sequence data are deposited in the DDBJ Sequence Read Archive (DRA Accession no. PRJDB16387).

**Bioinformatics**

Raw sequencing reads were demultiplexed using bcl2fastq v2.20 (Illumina), and the demultiplexed paired-end reads were merged using the `fastq_mergepairs` command in USEARCH v11.0.667 (Edgar 2010) with the default settings. Read quality filtering was performed using the USEARCH `fastq_filter` command with `-fastq_maxee 1.0 -fastq_minlen 180` options. cutPrimers v2.0 (Kechin *et al.* 2017) was used to remove the primer sequences, and the number of unique base sequences in preprocessed reads was calculated using the USEARCH `fastx_uniques` command. The USEARCH `unoise3` command with the default settings was used to perform error correction of amplicon reads. As a result, a set of predicted biological sequences called zero-radius operational taxonomic units (ZOTUs) was obtained. The USEARCH `otutab` command was used to map the quality-filtered reads in each sample to the generated ZOTUs at >97%

similarity. Finally, the highest number of ZOTU reads obtained from the negative controls for sampling, DNA extraction, and PCR was subtracted from the number of ZOTU reads in each sample.

For taxonomic assignments of the ZOTUs, `blastn` searches were performed of the Kanagawa Prefecture DNA database for aquatic insects (ver1.0(April 2023), `https://www.pref.kanagawa.jp/docs/b4f/suigen/edna-en.html`) using the local BLAST 2.13.0+ (Camacho *et al.* 2009) and NCBI GenBank database (Clark *et al.* 2016) using the online BLAST (accessed on March 2, 2023) with >99% identity. The former database is a collection of DNA sequences, mainly of aquatic insects found in Kanagawa Prefecture, which has been registered to promote research on aquatic insects using eDNA and adopts taxonomy conforming to the checklist for the National Census on River Environments of the Ministry of Land, Infrastructure, Transport and Tourism in Japan. Taxonomic assignments for each ZOTU were made up to the species level where possible using the results of the Kanagawa Prefecture DNA database as a priority. NCBI GenBank database results were used when they assigned a lower taxonomic level than the Kanagawa Prefecture DNA database. When only results up to the genus level were available for a given ZOTU, we considered them as valid and assigned that genus to the ZOTU. When two taxa had the same % identity for a given ZOTU, we considered them together as a single taxon.

In total, 247 taxa (henceforth referred to as ‘species’) of aquatic insects were detected, and 169 species that had occurred in four or more samples were modeled in the subsequent analysis. The degree of primer-template mismatches (the total number of mismatched bases in the priming region of the forward and reverse primers) of each species was determined by USEARCH `search_pcr` command.

The resulting dataset consists of 77 samples, as two samples from a site were lost because of filter clogging and one due to a DNA extraction error. The mean sequencing depth for the 77 samples was 40,416.6, with a standard deviation of 27,795.2.

### References

Camacho, C., Coulouris, G., Avagyan, V., Ma, N., Papadopoulos, J., Bealer, K. & Madden, T.L. (2009) Blast+: architecture and applications. *BMC Bioinformatics*, **10**, 421.

Clark, K., Karsch-Mizrachi, I., Lipman, D.J., Ostell, J. & Sayers, E.W. (2016) GenBank. *Nucleic Acids Research*, **44**, D67–D72.

Edgar, R.C. (2010) Search and clustering orders of magnitude faster than BLAST. *Bioinformatics*, **26**, 2460–2461.

Kechin, A., Boyarskikh, U., Kel, A. & Filipenko, M. (2017) cutPrimers: a new tool for accurate cutting of primers from reads of targeted next generation sequencing. *Journal of Computational Biology*, **24**, 1138–1143.

O'Donnell, J.L., Kelly, R.P., Lowell, N.C. & Port, J.A. (2016) Indexed PCR primers induce template-specific bias in large-scale DNA sequencing studies. *PLoS ONE*, **11**, e0148698.

Takenaka, M., Hasebe, Y., Yano, K., Okamoto, S., Tojo, K., Seki, M., Sekiguchi, S., Jitsumasa, T., Morohashi, N., Handa, Y. & Sakaba, T. (2024) Environmental DNA metabarcoding on aquatic insects: comparing the primer sets of MtInsects-16S based on the mtDNA 16S and general marker based on the mtDNA COI region. *Environmental DNA*, **6**, e588.

Takenaka, M., Yano, K., Suzuki, T. & Tojo, K. (2023) Development of novel PCR primer sets for DNA barcoding of aquatic insects, and the discovery of some cryptic species. *Limnology*, **24**, 121–136.

Wong, M.K.S., Nakao, M. & Hyodo, S. (2020) Field application of an improved protocol for environmental DNA extraction, purification, and measurement using Sterivex filter. *Scientific Reports*, **10**, 21531.
